# Assessing the Accuracy of a Smart Collar for Dogs: Predictive Performance for Heart and Breathing Rates on a Large Scale Dataset

**DOI:** 10.1101/2023.06.09.544347

**Authors:** Hugo Jarkoff, Guillaume Lorre, Eric Humbert

## Abstract

We present an evaluation of a smart collar for dogs that measures resting heart and breathing rates. The study involved 40 dogs of various breeds, ages, and sizes. The collar’s accuracy in estimating heart rates, breathing rates and detecting heart pulses was evaluated by comparing the measurements from the collar with those from a portable ECG device, completed by labels annotated by trained humans. The results show that the collar provides accurate and reliable measurements of heart and breathing rates, with a (symmetric) mean absolute percentage error of 0.38% and 1.42% respectively, and a F1 score of 98.04% for the detection of heart beats at 50ms precision.

Our study demonstrates the potential of Invoxia’s *Smart Dog Collar* as a medical-grade tool for continuous, noninvasive, remote monitoring of vital signs in dogs, helping to improve the quality of life for dogs, detect early signs of illness, and strengthen the bond between dogs and their owners.

## 1 Introduction

Wearable technology has become increasingly important for monitoring human health, with devices employing miniature optical sensors worn on the wrist or finger to measure cardiac activity. Such information could undoubtedly improve the health of our canine companions as well. Indeed, several studies have shown that dogs can suffer from a wide range of health issues, including cardiovascular disease, respiratory disease, and cancer. Such conditions can be linked to changes in resting heart and breathing rates, which are important indicators of health and well-being in dogs (Porciello et al., 2016; Rishniw et al., 2012; Schober et al., 2011). However, due to their fur, non-invasive optical or electrical sensors are not feasible for use on dogs.

In response, Invoxia has developed an innovative solution incorporated into a lightweight wireless dog collar, the *Smart Dog Collar*, which utilizes ultra-precise motion sensors and innovative artificial intelligence algorithms to non-invasively measure a dog’s resting heart and respiratory rates throughout the day and night. A depiction of the collar is shown in Figure 1. The method also detects heart beats leading to the estimation of RR intervals (time elapsed between two successive R waves of the QRS signal on an electrocardiogram). This makes it possible to analyze heart arrhythmia and variability, for instance through a Poincaré plot.

**Figure 1.**
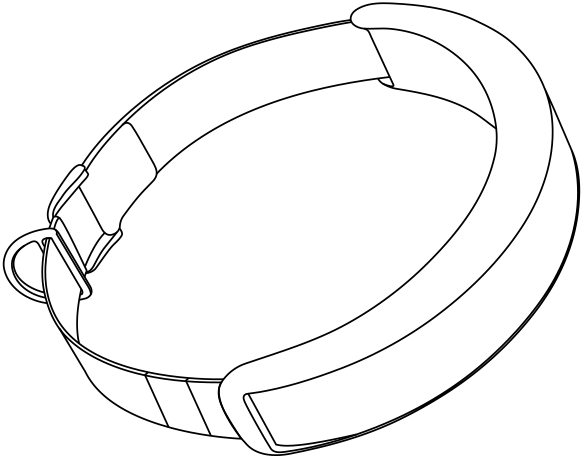
Depiction of the *Smart Dog Collar* from Invoxia. The collar is lightweight (165g including the strap) and measures 175 × 88 × 34mm. See smartdogcollar.com.

Several studies have been conducted to evaluate the accuracy of similar wearable devices for measuring heart and breathing rates in dogs. However, these studies either featured a small number of dogs (Foster et al., 2020a, Foster et al., 2020b, Brugarolas et al., 2015, Lahdenoja et al., 2019), or were conducted in a laboratory setting (Essner et al., 2015), or used a bulky harness that is not suitable for long-term continuous monitoring of dogs (Foster et al., 2020a, Brugarolas et al., 2015). All of these results featured a large variability in the accuracy of the devices. To the best of our knowledge, the only study that evaluated a similar collar for dogs was conducted by Foster et al., 2020b. The authors reported an an average accuracy of heart rate estimation of 88.2%, and an average accuracy of respiration rate estimation of 94.3%. However, the study was conducted on a single dog, with a very small number of measurements (19 minutes of data collection), which can be insufficient to evaluate the accuracy of the algorithms.

The goal of this article is to present a large-scale study evaluating the metrics estimated by Invoxia’s *Smart Dog Collar*, specifically the estimations of resting heart and respiratory rates, as well as the detection of heartbeats. To achieve this, we will first introduce the evaluation dataset, followed by the definitions of the metrics used to measure the quality of the algorithms. Then, we will discuss our machine learning models training and evaluation methodology. Finally, we will present and discuss the results obtained.

## 2 Dataset

The machine learning algorithms used in this study are trained using k-fold cross-validation on a training dataset and then tested on an independent test dataset ^1^. In this section, we will provide a comprehensive description of the test dataset utilized for evaluating the performance of the *Smart Dog Collar*’s algorithms.

### 2.1 Dog Participants

Our test dataset comprises data from 40 dogs, representing a diverse range of breeds, ages, and weights. The participating dogs were selected to ensure a wide variety of characteristics, providing a robust dataset for algorithm evaluation. The dataset includes small, medium, and large dog breeds, with ages ranging from young to senior dogs. Their fur also range from thin to thick.

### 2.2 Data Collection

The data collection process involved fitting each dog with a *Smart Dog Collar*, which records the signals from its integrated motion sensors. Simultaneously, a portable ECG device was attached to the dog and connected via Bluetooth to the *Smart Dog Collar*. The ECG device transmitted and recorded ECG signals sampled at 120 Hz, which were then stored by the collar. Each recording session took place while the dog was sleeping, and lasted on average 300s, with multiple sessions per day. The duration of the recordings ensured that sufficient data was collected for algorithm evaluation, while also minimizing potential discomfort for the dogs. The total duration of the evaluation recordings was 5585 minutes (93 hours). However, some of the recording sessions were discarded due to poor signal quality (too many movements from the dog), which resulted in some dogs having less than 300 seconds of data available for evaluation. All recordings sessions were then divided into smaller segments (or *samples*, as referred later in this article) for evaluation by our algorithms. All recordings sessions took place in the dog’s home environment, to ensure that the dogs were comfortable and relaxed (which may not be the case in a veterinary clinic or laboratory setting), thus providing a realistic dataset for algorithm evaluation. To the best of our knowledge, this is the first study to evaluate a wearable device for dogs in a relaxed home environment, on a large scale, and for a long period of time.

### 2.3 Annotation

To obtain accurate ground truth data for the resting heart and respiratory rates, each recording session was filmed using a high-definition video camera. The video footage was then synchronized with the recorded sensor and ECG signals, providing a comprehensive dataset for further analysis. The synchronized video footage was annotated by trained experts to extract the respiratory rate over time. The annotators carefully observed the dog’s chest movements and counted the number of breaths per minute throughout the recording session. The ECG recordings were also semi-automatically annotated using custom software, which identified the exact instances of heartbeats (R-peaks) within the ECG signal. The software’s output was then manually reviewed and adjusted by experts, ensuring the accuracy of the detected heartbeats. Each recording session was also preprocessed using a custom algorithm to filter out the movements of the dog and the noise from the ECG signal. Thus, only the resting heart and respiratory rates were extracted from the data.

### 2.4 Dataset Characteristics

The different (anonymized) dogs constituting our test dataset are shown in the table 1. The table includes the dog’s ID, its main breed, its weight in kilograms, and its age in years. Some descriptive figures of the dataset are also provided in the figures 2a, 2b, 2c and 3. We can see that our test dataset consists mainly of relatively young, small to mid-sized dogs. Based on the different breeds, the furs range from very thin (whippet) to extremely thick (samoyede). Therefore, our test dataset can be thought as a solid foundation for evaluating the performance of the *Smart Dog Collar*’s algorithms in estimating resting heart and respiratory rates.

**Table 1.** Dog participants

| **Id** | **Breed (main)** | **Weight (kg)** | **Age (years)** |
| --- | --- | --- | --- |
| 0 | beagle | 9.3 | 4.5 |
| 1 | beagle | 10.0 | 4.5 |
| 2 | beauceron | 29.0 | 7.5 |
| 3 | beauceron | 38.0 | 4.0 |
| 4 | australian_shepherd | 20.0 | 9.5 |
| 5 | braque_de_weimar | 27.0 | 2.5 |
| 6 | cocker_anglais | 15.4 | 2.5 |
| 7 | unknown | 18.2 | 2.5 |
| 8 | unknown | 17.5 | 2.0 |
| 9 | unknown | 23.5 | 2.5 |
| 10 | basset_fauve_de_bretagne | 16.0 | 2.5 |
| 11 | berger | 11.0 | 5.5 |
| 12 | berger | 10.5 | 4.0 |
| 13 | australian_shepherd | 26.0 | 2.0 |
| 14 | boxer | 34.0 | 4.5 |
| 15 | weimaraner | 27.0 | 5.5 |
| 16 | dalmatien | 17.0 | 2.0 |
| 17 | epagneul | 14.2 | 3.0 |
| 18 | epagneul | 15.0 | 3.5 |
| 19 | leonberg | 42.0 | 1.5 |
| 20 | setter | 24.0 | 3.0 |
| 21 | dalmatien | 24.0 | 2.5 |
| 22 | dalmatien | 25.0 | 1.0 |
| 23 | doberman | 33.0 | 1.5 |
| 24 | fox_terrier | 9.0 | 9.5 |
| 25 | husky | 29.0 | 4.0 |
| 26 | labrador | 25.0 | 3.5 |
| 27 | labrador | 28.0 | 1.5 |
| 28 | labrador | 27.0 | 4.0 |
| 29 | staffie | 19.0 | 5.5 |
| 30 | tervueren | 24.8 | 2.0 |
| 31 | berger_blanc_suisse | 29.0 | 1.5 |
| 32 | unknown | 20.0 | 5.5 |
| 33 | samoyede | 25.0 | 3.5 |
| 34 | golden_retriever | 22.0 | 1.1 |
| 35 | golden_retriever | 22.0 | 1.1 |
| 36 | whippet | 11.7 | 2.5 |
| 37 | labrador | 30.0 | 1.2 |
| 38 | australian_shepherd | 23.0 | 2.1 |
| 39 | dogo_argentino | 30.0 | 6.1 |

**Figure 2.**
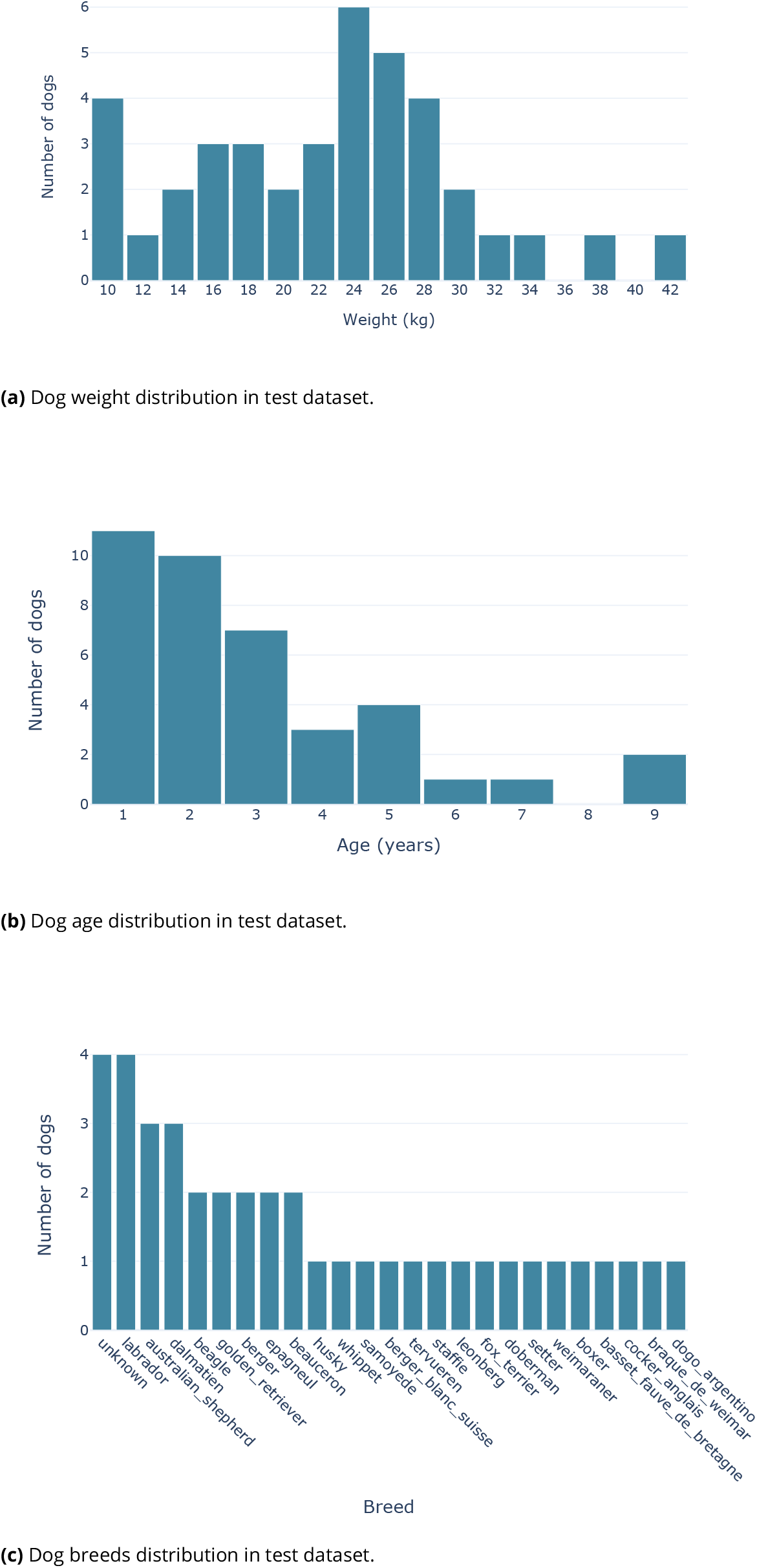
Test dataset characteristics (age, weight, and breeds).

**Figure 3.**
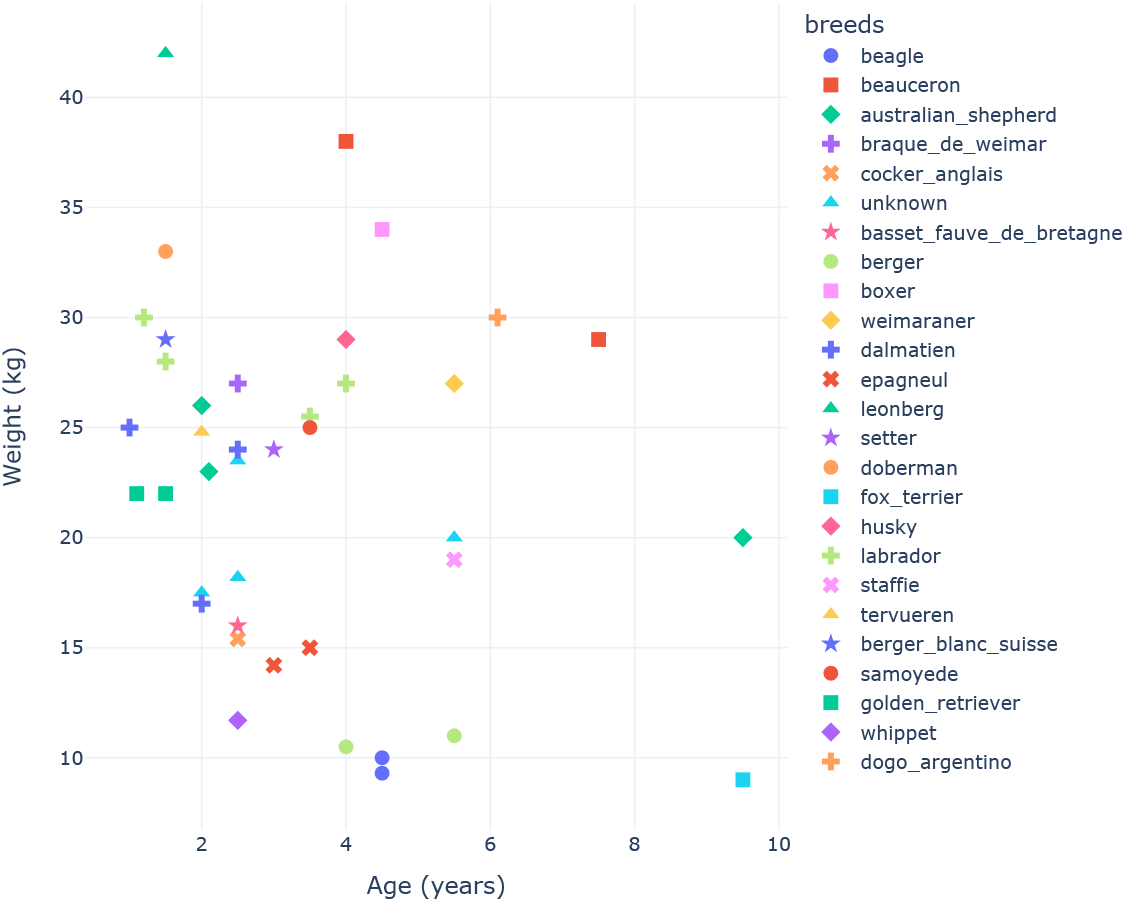
Distribution of dogs weight vs age in test dataset, with breeds as colors.

## 3 Evaluation Metrics

To assess the performance of the *Smart Dog Collar*’s algorithms, it is essential to establish a set of evaluation metrics that allow for a quantitative comparison between the collar’s estimations and the ground truth data obtained from the annotated dataset. In this section, we will introduce and describe the chosen metrics for evaluating the collar’s ability to estimate heart and respiration rates, as well as its effectiveness in detecting pulses.

### 3.1 Heart Rate Error and Heart Rate Accuracy

A popular metric to use in regression problem is the Mean Absolute Percentage Error (MAPE), defined by :

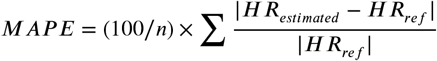

Where *HR*_*ref*_ (*resp. HR*_*estimated*_) represents the Heart Rate of reference (*resp*. estimated by the machine learning algorithm) and *n* is the number of samples. An issue with MAPE is that it is not bounded, meaning that the point-wise error can be arbitrarily large.

Therefore, in our study, we found the Symmetric Mean Absolute Percentage Error (SMAPE) to be more suitable as an evaluation metric for our machine learning algorithm. SMAPE is defined by :

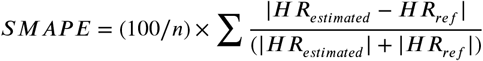

SMAPE is a widely used metric in regression problems (Törnqvist, Vartia, and Vartia, 1985), which measures the average of the absolute error (distance) between the prediction and label, over the sum of the absolute values of the prediction and label. Rewritten as :

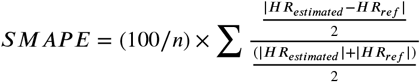

SMAPE computes the (average of the) halved distance between the prediction and label, over the midpoint of the prediction and label. SMAPE has a bounded range of [0, 100], where 0 indicates perfect accuracy and 100 indicates complete inaccuracy. This range is more intuitive than the range of MAPE, which is unbounded and difficult to interpret.

From SMAPE, we defined the Symmetric Mean Absolute Percent Accuracy (SMAPA), which measures, for each sample, the relative accuracy between the estimated heart rate provided by the *Smart Dog Collar* and the reference heart rate obtained from the annotated ECG data:

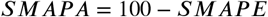

SMAPA provides an intuitive measure of how close the collar’s estimations are to the true heart rate values, with values closer to 100 indicating better performance. As these measures essentially represent the same information, we will use both SMAPE and SMAPA interchangeably in the rest of this paper. Other measures are also used in the literature to evaluate the performance of heart rate estimation algorithms, such as “readout error of no greater than ±10% of the input rate or ±5 beats per minute (bpm), whichever is greater” (Medical Instrumentation et al., 2002, Jacobs et al., 2021, Li et al., 2019) as defined by the American National Standards Institute standard for cardiac monitors, heart rate meters, and alarms. However, we found that these measures are not suitable for our study, as they are not sensitive enough to evaluate the performance of the *Smart Dog Collar*.

### 3.2 Pulse Detection Precision and Pulse Detection Recall

In order to determine whether a pulse is considered detected, we compare the timing of the estimated pulse provided by the *Smart Dog Collar* with the reference pulse obtained from the annotated ECG data. A pulse is considered detected if it occurs within a specified time window around the reference pulse:

- A pulse occurring within ±50 ms of the reference pulse is considered very good according to the literature (Blok et al., 2021) ;
- A pulse occurring within ±100 ms of the reference pulse can still be acceptable, especially for dogs as they have a larger variability of RR intervals than humans.

By using these criteria, we can evaluate the collar’s effectiveness in detecting heartbeats in terms of precision and recall, providing a comprehensive understanding of its performance in canine health monitoring.

#### Pulse Detection Precision

measures the proportion of relevant pulses (correctly detected heartbeats) among the total retrieved pulses (all detected heartbeats). It is calculated as:

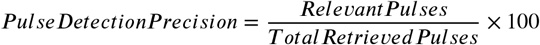

#### Pulse Detection Recall

measures the proportion of relevant pulses (correctly detected heartbeats) among the total pulses present in the dataset (all true heartbeats). It is calculated as:

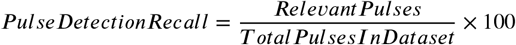

These two metrics together provide a comprehensive evaluation of the collar’s effectiveness in detecting heartbeats, taking into account both false positives (incorrectly detected heartbeats) and false negatives (missed heartbeats). These two metrics can be combined by harmonic mean defining the F1 score :

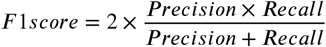

It is widely used in machine learning to evaluate the performance of detection algorithms as it provides a balanced measure of the two metrics. The F1 score is bounded between 0 and 1, and higher values indicate better performance. A high F1 score indicates high precision and high recall.

### 3.3 Respiration Rate Error and Respiration Rate Accuracy

We also evaluate the collar’s performance in estimating respiration rates using the same metrics as for heart rate :

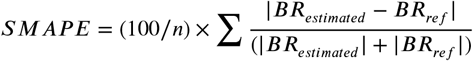

Where *BR*_*ref*_ represents the Breathing Rate of reference obtained from the annotated video footage, and *BR*_*estimated*_ represents the Breathing Rate estimated by the *Smart Dog Collar*’s machine learning algorithm and *n* is the number of samples. SMAPA is also rigorously similar to the one defined for heart rate, and derived from SMAPE in the same way. These metrics allow for a quantitative evaluation of the collar’s performance in estimating respiration rates.

By employing these metrics, we can rigorously assess the performance of the *Smart Dog Collar*’s algorithms in estimating resting heart and respiratory rates and detecting heartbeats, enabling a thorough understanding of the device’s capabilities and potential for use in canine health monitoring.

## 4 Training methodology

We provide details about the training methodology used to train and evaluate our machine learning models.

### 4.1 K-Folds cross-validation

We use regular K-folds cross-validation (see figure 4) to assess our machine learning algorithm training protocol. K-fold cross-validation is a statistical method used to evaluate the performance of machine learning models on a limited amount of data. In this method, our original dataset is divided into K equally sized subsets, or *folds*. One of these folds is retained as the validation set, and the remaining K-1 folds are used as the training set. Our machine learning model is then trained on the training set and evaluated on the validation set. This process is repeated K times, with each fold serving as the validation set exactly once. This validation performance is used to tune the hyperparameters of the model and detect error in labels without using the test set. Only the final models (after hyperparameter tuning) are evaluated on the test set, and the results of these K evaluations over the test set are then averaged to produce a single estimation of the model’s performance. This procedure allow us to assess the model’s generalization performance on unseen data (the test set) and avoid overfitting on the test set.

**Figure 4.**
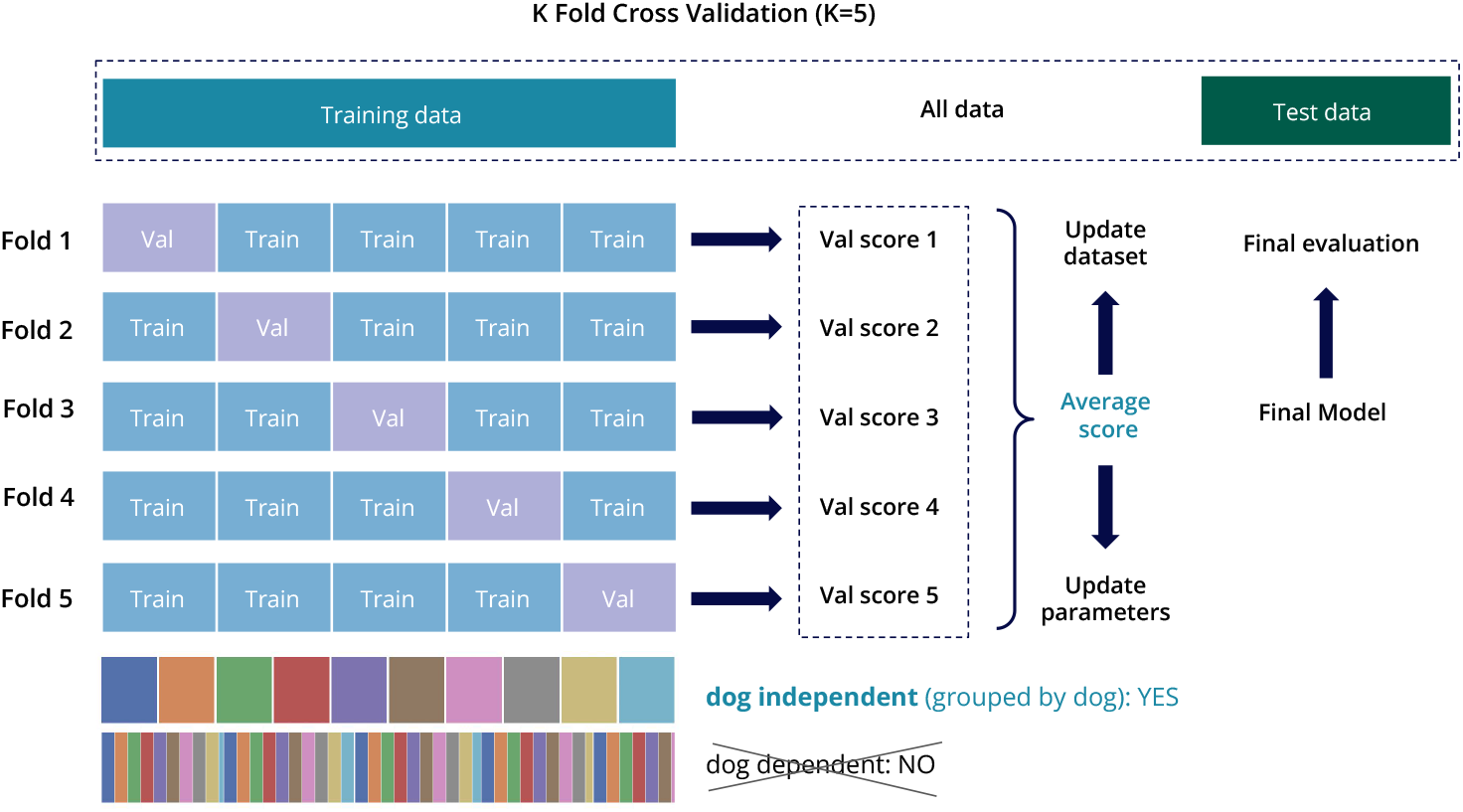
K-Folds cross-validation procedure description. The number of folds depicted (5) is not be representative of the actual number of folds used.

### 4.2 Data preprocessing

When evaluating our trained machine learning models, we must first preprocess the data to remove all noises and artifacts. Indeed, while our data was collected in a controlled environment, with the dogs being at rest, small agitations and movements from the dogs can still occur (*e*.*g*, dreaming, changing rest position…). These movements can be captured by either the ECG device or the collar and cause artifacts in the data, essentially hiding the heart and respiratory micromovements, which can be problematic for the evaluation of our machine learning models. Therefore, we must remove these artifacts from the data before evaluating our models. To do so, we use a custom preprocessing algorithm that identifies and removes such artifacts from the data. After applying this algorithm, we remove 16.6% of the total data ^2^, which is a reasonable amount. Upon visual inspection, we found that the remaining data is of high quality, with no visible artifacts. This clean dataset is then used to evaluate our models using the metrics described in section 3.

## 5 Results and discussion

Once our Heart and Breathing rates algorithms are trained using K-folds cross-validation, we evaluate their respective performance on the test set using the metrics described in section 3. We also evaluate the performance of our Heartbeat detection algorithm using the metrics described in the previous section.

### 5.1 Heart Rate results

*The Smart Dog Collar* makes it possible to predict the timing of the cardiac pulsations and estimate the heart rate. Comparing it with the reference heart pulses obtained by ECG, it is possible to evaluate the performance of the model with the metrics described in the section 3. The model is trained K times with K-folds cross-validation and table 2 summarizes the metrics obtained. The best, worst and the mean of the K models are presented. The results on the different folds are very similar with a standard deviation inferior to 0.1% for all metrics. Therefore, training has been done on a large quantity of data that prevents the model from overfitting. The mean SMAPE is very low (0.38%) showing that the model is able to estimate the heart rate with a very high precision. The timing of the cardiac pulsations is also most of the time predicted within a margin of error of 50ms, given the f1score (50ms) of 97.93%. This precision in the timing of cardiac pulsations is necessary to study the arrhythmias and the heart rate variability. Furthermore, very few pulses are missed or incorrectly detected, with a recall of 99.24% and a precision of 98.96% at 100ms.

**Table 2.** Heart pulse detection and heart rate estimation results on test set. Cross validation gives us K models, the results from the worst and best models are presented as well as the mean and standard deviation.

| Folds statistics | SMAPE | f1score |  | precision |  | recall |  |
| --- | --- | --- | --- | --- | --- | --- | --- |
|  |  | 50 ms | 100 ms | 50 ms | 100 ms | 50 ms | 100 ms |
| min | 0.37 | 97.95 | 99.04 | 97.82 | 98.91 | 98.11 | 99.21 |
| max | 0.40 | 98.14 | 99.12 | 98.01 | 99.01 | 98.30 | 99.28 |
| mean | 0.38 | 98.04 | 99.08 | 97.93 | 98.96 | 98.20 | 99.24 |
| std | 0.01 | 0.08 | 0.04 | 0.08 | 0.04 | 0.08 | 0.03 |

The table 3 details the distribution of the metrics across all the samples. Half of the samples have perfect predictions with no error. Only 5% of the examples have a SMAPA inferior to 97%. The collar is consequently able to detect heart beats quite effectively even in the worst cases.

**Table 3.** Heart pulse detection and heart rate estimation results on test set with percentiles results across all samples.

|  | SMAPE | f1score |  | precision |  | recall |  |
| --- | --- | --- | --- | --- | --- | --- | --- |
|  |  | 50 ms | 100 ms | 50 ms | 100 ms | 50 ms | 100 ms |
| mean | 99.62 | 98.04 | 99.08 | 97.93 | 98.96 | 98.20 | 99.24 |
| std | 1.22 | 4.96 | 2.96 | 5.28 | 3.45 | 4.84 | 2.83 |
| 5% | 97.00 | 88.90 | 93.80 | 88.14 | 92.90 | 88.90 | 93.80 |
| 25% | 99.93 | 100.00 | 100.00 | 100.00 | 100.00 | 100.00 | 100.00 |
| 50% | 99.97 | 100.00 | 100.00 | 100.00 | 100.00 | 100.00 | 100.00 |
| 75% | 99.97 | 100.00 | 100.00 | 100.00 | 100.00 | 100.00 | 100.00 |
| 95% | 100.00 | 100.00 | 100.00 | 100.00 | 100.00 | 100.00 | 100.00 |

Next, the variability of the performance of the model is studied across the different breeds, age and sizes of dogs as well as across the heart rate distribution. The results are quite similar across all breeds as presented in the figure 5a. The worst breed has a SMAPE of 1.7% (still quite low) but this breed has also few examples which is not representative. The performance does not vary much across the different ages and the different weights of the dog (5b and 5c). The worst weight bin and worst age bin have a very low SMAPES (1.25% and 1.17% respectively). The collar is therefore suitable for all dogs regardless of their age, weight or breed. Even if the model has bigger errors on the extremities of the heart rate distribution, it is still acceptable (inferior to 2% SMAPE) (see figure 6a). Another point is that these are quite rare cases and therefore less represented in the training set. The collar should therefore be able to monitor the heart rate even if it is out of the normal range.

**Figure 5.**
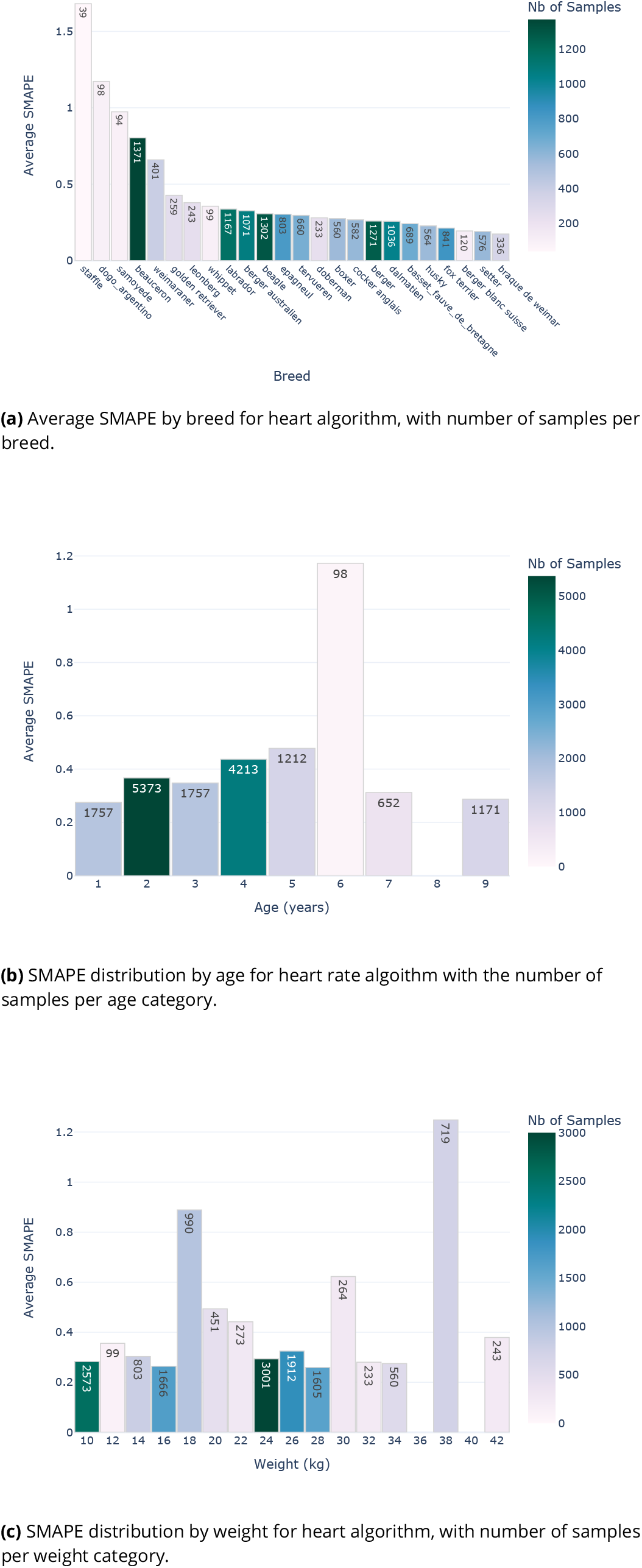
SMAPE distribution for different dog characteristics (breed, age and weight) for heart rate estimation algorithm.

**Figure 6.**
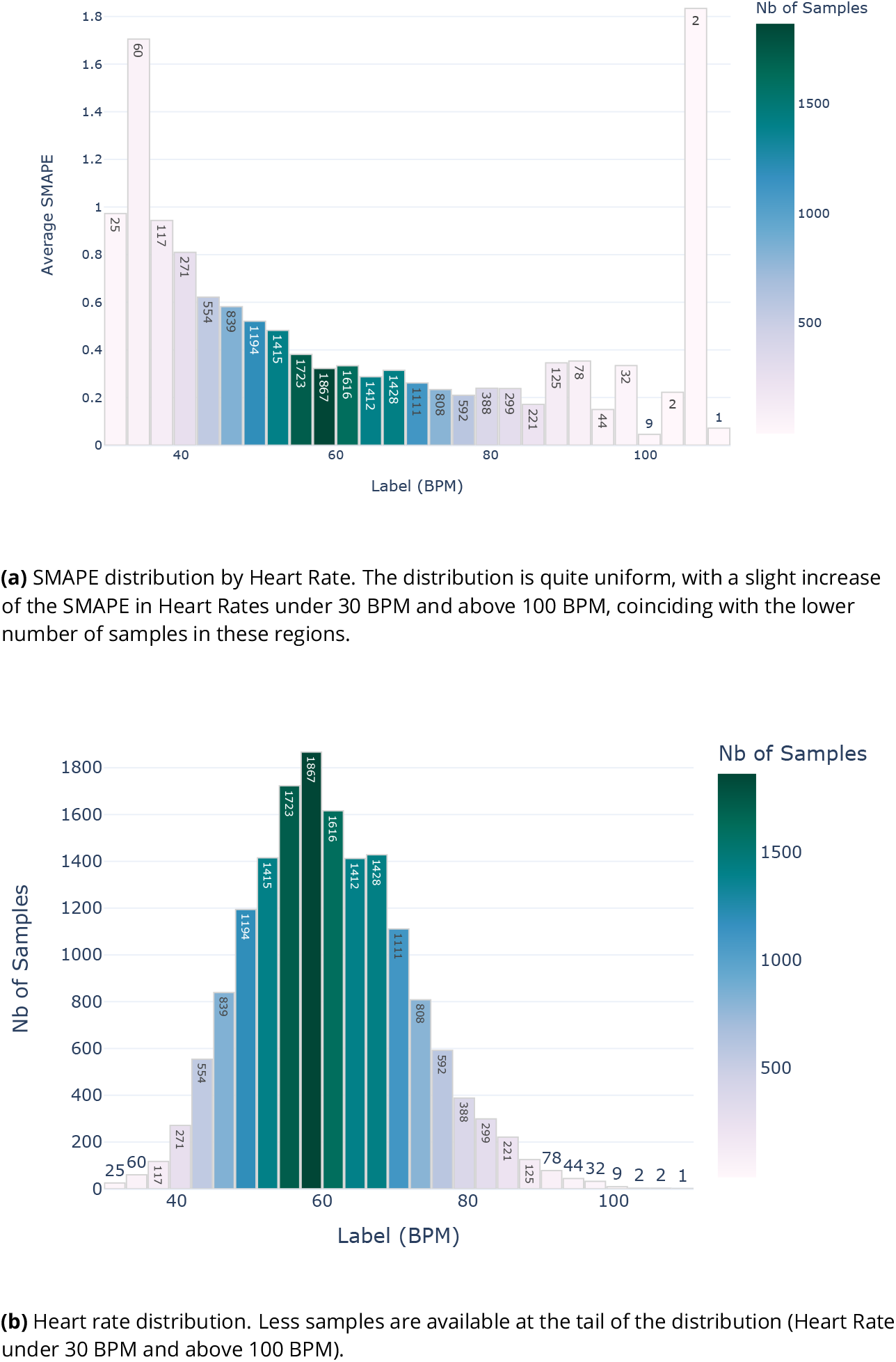
SMAPE and Label distribution by ground truth for Heart Rate estimation algorithm.

**Figure 7.**
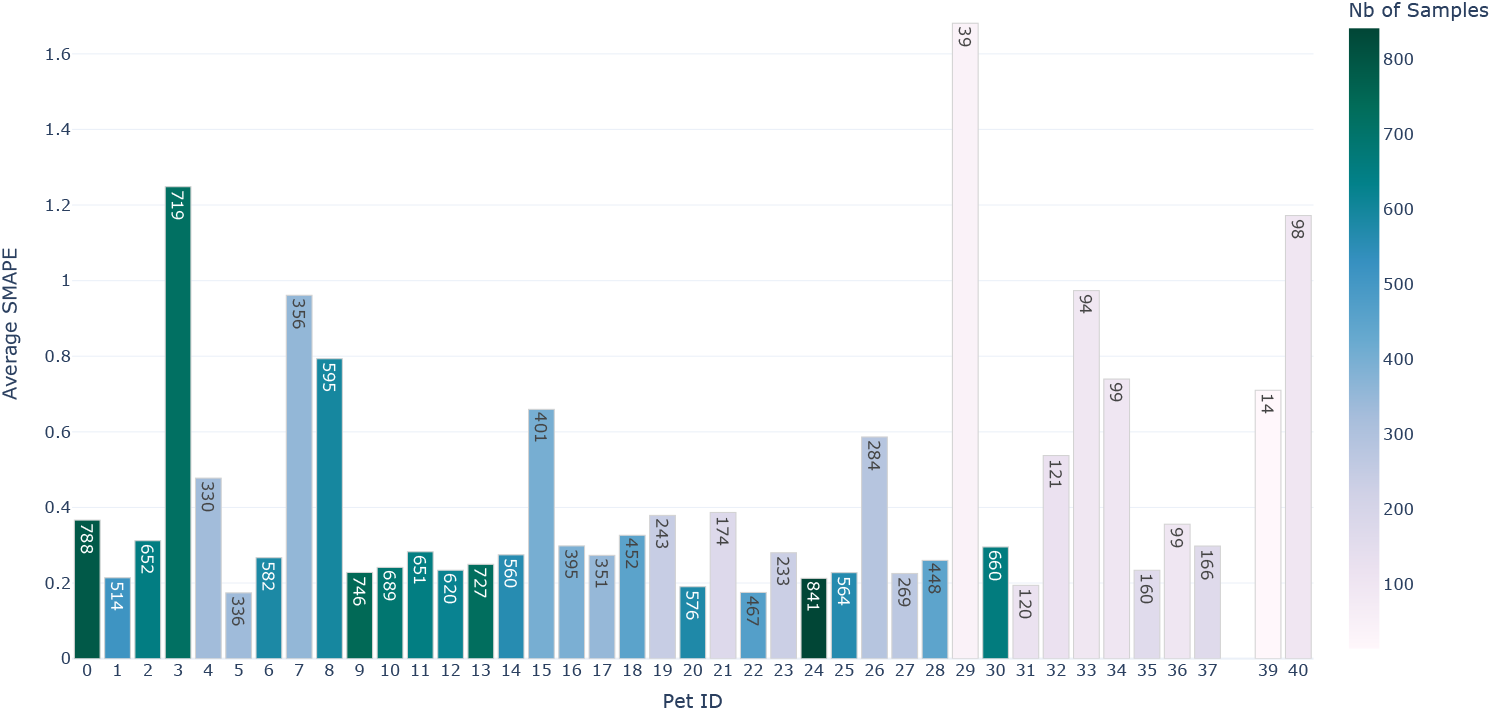
Heart SMAPE for all the dogs in the test set. The color represents the quantity of samples.

Finally, we give a more visual understanding of the performance of the model. Figure 8 shows the predicted heart rate against the ground truth for every sample in the test data. Transparency is used to better show the density of points. Most of the points are on the diagonal which means that predictions are very close to the ground truth. Lines above and below the diagonal can be explained by a missing or extra beat in the prediction. This figure also highlights the fact that the model is good in all the range of heart rate.

**Figure 8.**
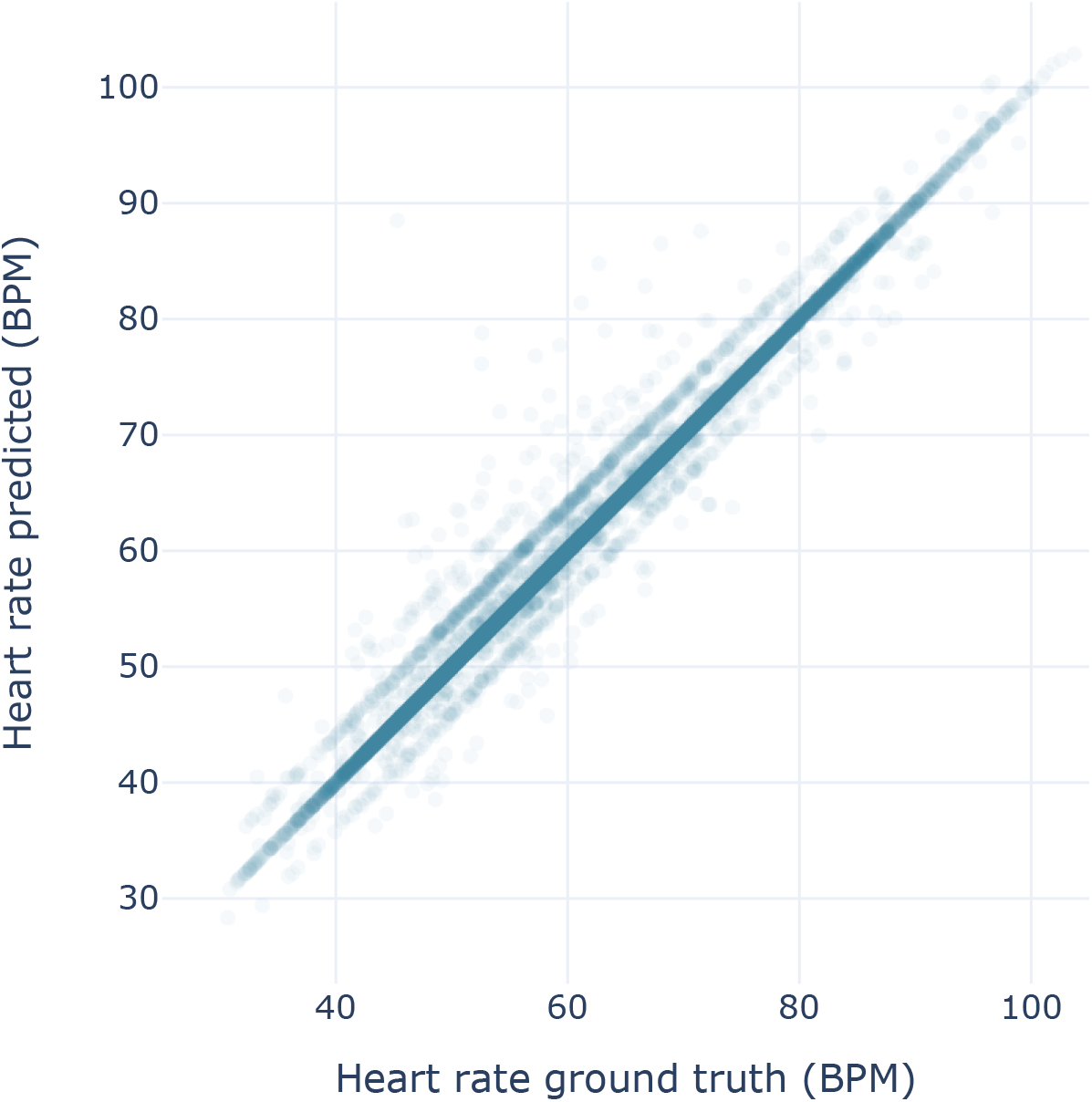
Comparison of the predicted heart rate and the ground truth. Transparency is used to better show the density of points. The diagonal represents perfect prediction.

On an individual level, figure 9 shows the distribution of the heart rate by dog for the prediction and the ground truth. The distributions of heart rate varies a lot across dogs but the model is able to estimate it correctly except for small outlier points. The predictions of our algorithm in conjonction with the ECG are shown in figure 10. The pulses predicted correspond well with the R-peaks of the ECG.

**Figure 9.**
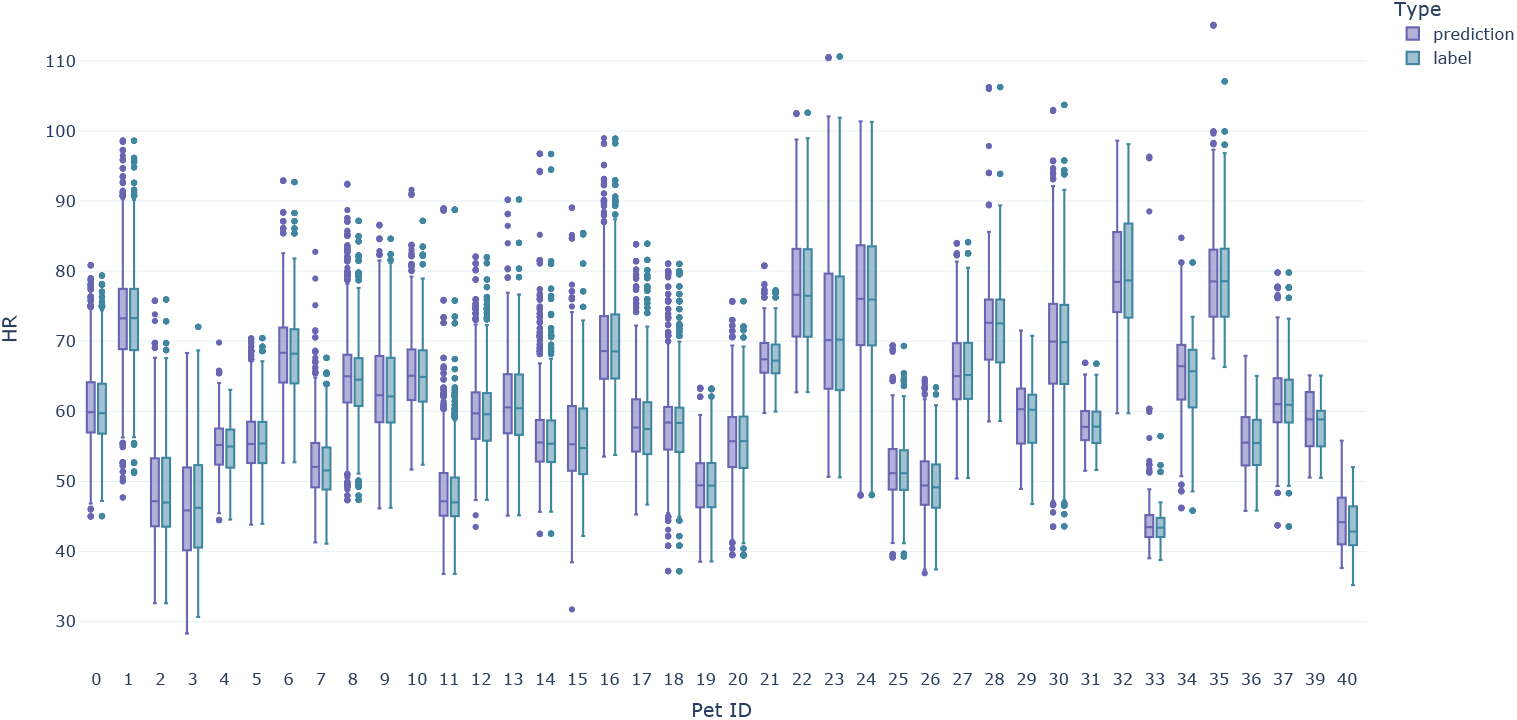
Heart rate distribution (ground truth vs predicted) by dog. The box plot shows the median, the first and third quartile and the outliers.

**Figure 10.**
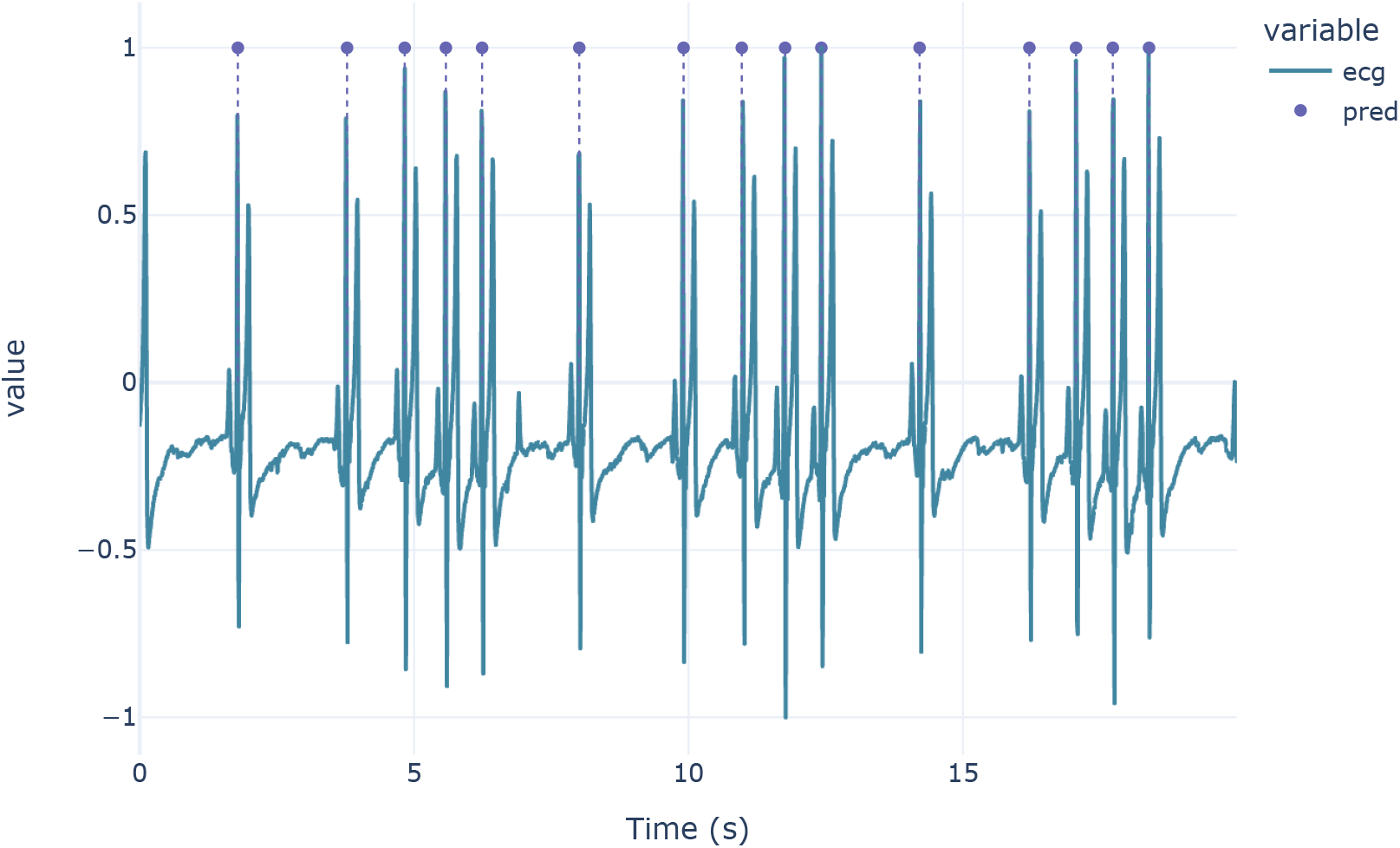
Comparison between the pulses predicted by our model and the ecg. The pulses are represented by dots.

To validate that our model is precise enough to study the heart rate variability and arrhythmias, the Poincaré plots with ground truth and labels are compared in figure 11. The Poincaré plot is a scatter plot of the RR intervals against the previous RR intervals. It is a common representation used in medical field, and is used to study the heart rate variability and arrhythmias and identify potential treatments (Moïse, Flanders, and Pariaut, 2020). The Poincaré plots obtained with the predicted pulses are very similar to the one obtained with the ground truth, which implies that our device is capable of producing medical-grade Poincaré plot useful for the study of heart rate variability and arrhythmias.

**Figure 11.**
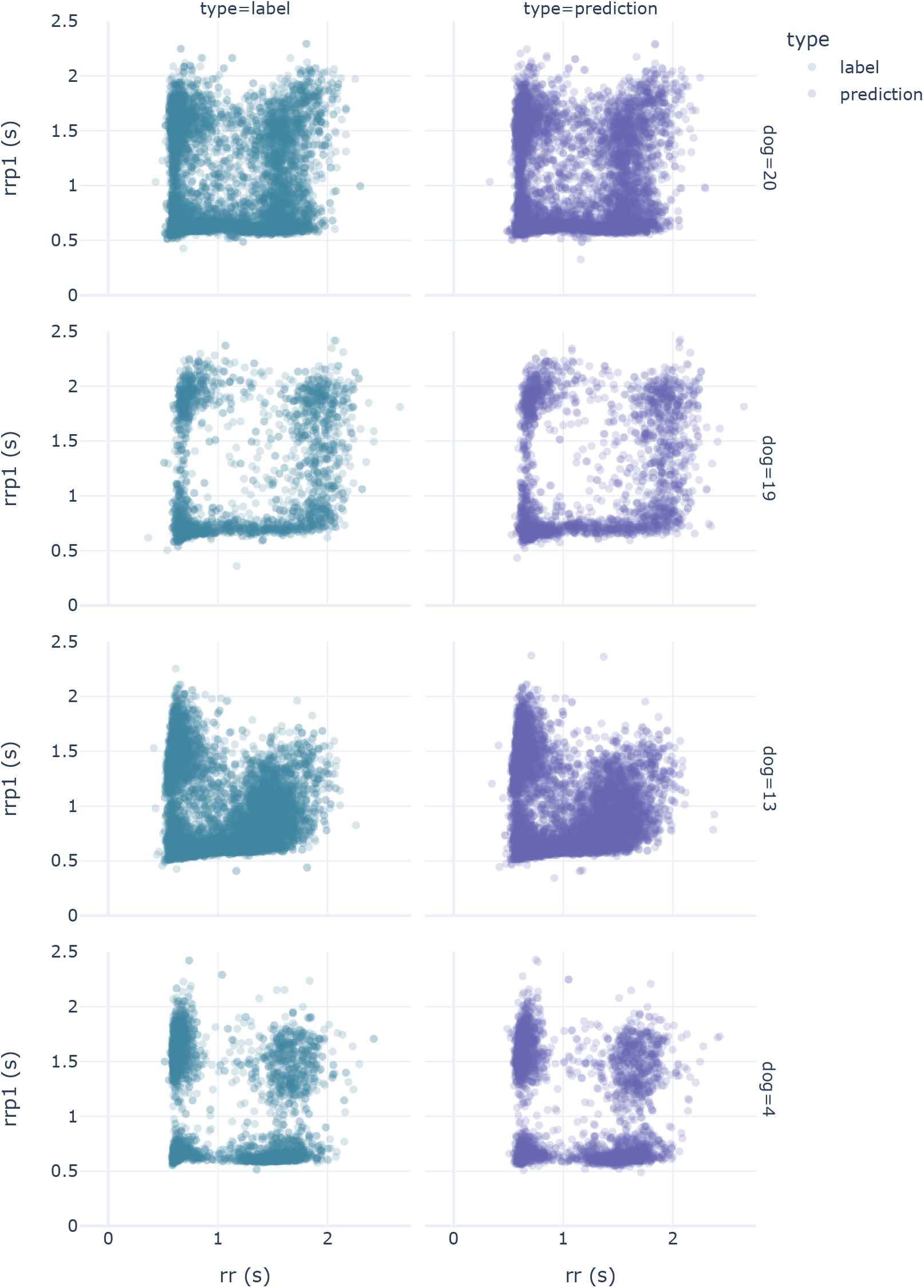
Comparison between the Poincaré diagram obtained with ground truth and predicted pulses.

### 5.2 Respiration Rate results

Similarly to the heart rate estimation, we first present the results of the respiration rate estimation on the test set. The model was trained on K folds of the training set, and the aggregated results over all folds are presented in the table 4.

**Table 4.** Breathing rate estimation results on test set. Cross validation gives us K models, the results from the worst and best models are presented as well as the mean and standard deviation.

| Folds statistics | SMAPE |
| --- | --- |
| min | 1.40 |
| max | 1.48 |
| mean | 1.42 |
| std | 0.04 |

The mean SMAPE over all folds is quite low (1.42%), and the standard deviation is also very low (0.04%). These results show that the model has a good generalization performance, and is not overfitting the training set. The table 5 details the distribution of the metrics across all the samples.

**Table 5.** Breathing rate estimation results on test set with percentiles results across all examples.

| Samples statistics | SMAPE |
| --- | --- |
| mean | 1.42 |
| std | 2.13 |
| 5% | 0.08 |
| 25% | 0.38 |
| 50% | 0.85 |
| 75% | 1.65 |
| 95% | 4.55 |

The results are still very good, with a mean SMAPE of 1.42% across all samples, and a standard deviation of 2.13%. The percentiles results show that 95% of the samples have a SMAPE below 4.55%, and 75% of the samples have a SMAPE below 1.65%. These results suggest that the model is able to generalize well on all samples, and could be used to estimate the breathing rate of any dog on a daily basis. To support these results, we study the variability of the performance of the model across the different dogs, and against their respective age, breed and size.

The results regarding individual dogs are presented in figure 12. All dogs have a SMAPE below 3%, which is a very good result. The distribution is fairly uniform, with some dogs having a SMAPE below 1% and some dogs having a SMAPE above 2%. These results show that the model is able to generalize well on all unseen dogs, regardless of their age, breed or size. To further support this claim, we present the results of the model against their respective breed, age and size on figures 13a, 13b and 13c.

**Figure 12.**
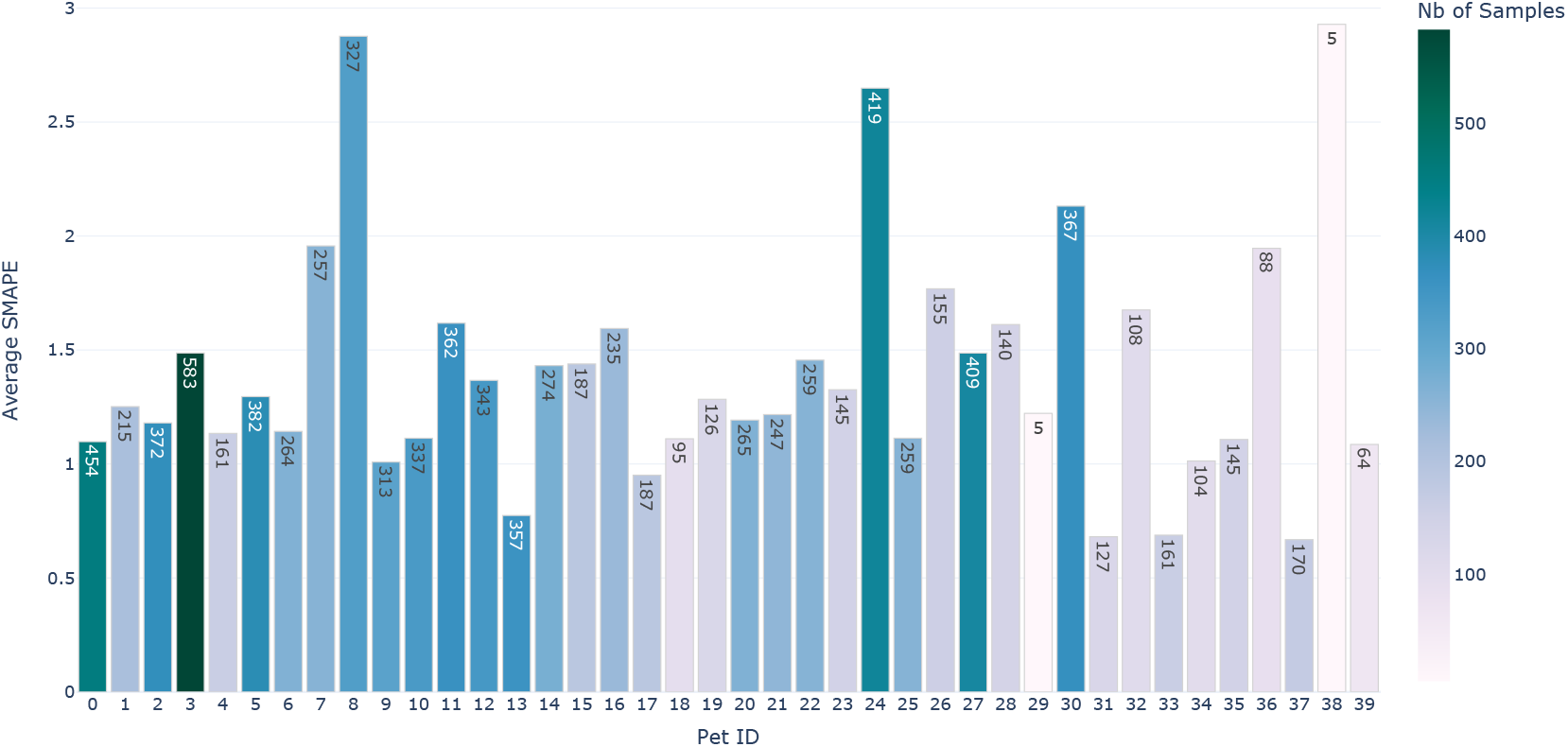
Average SMAPE by dog for breathing algorithm, with number of samples per dog.

**Figure 13.**
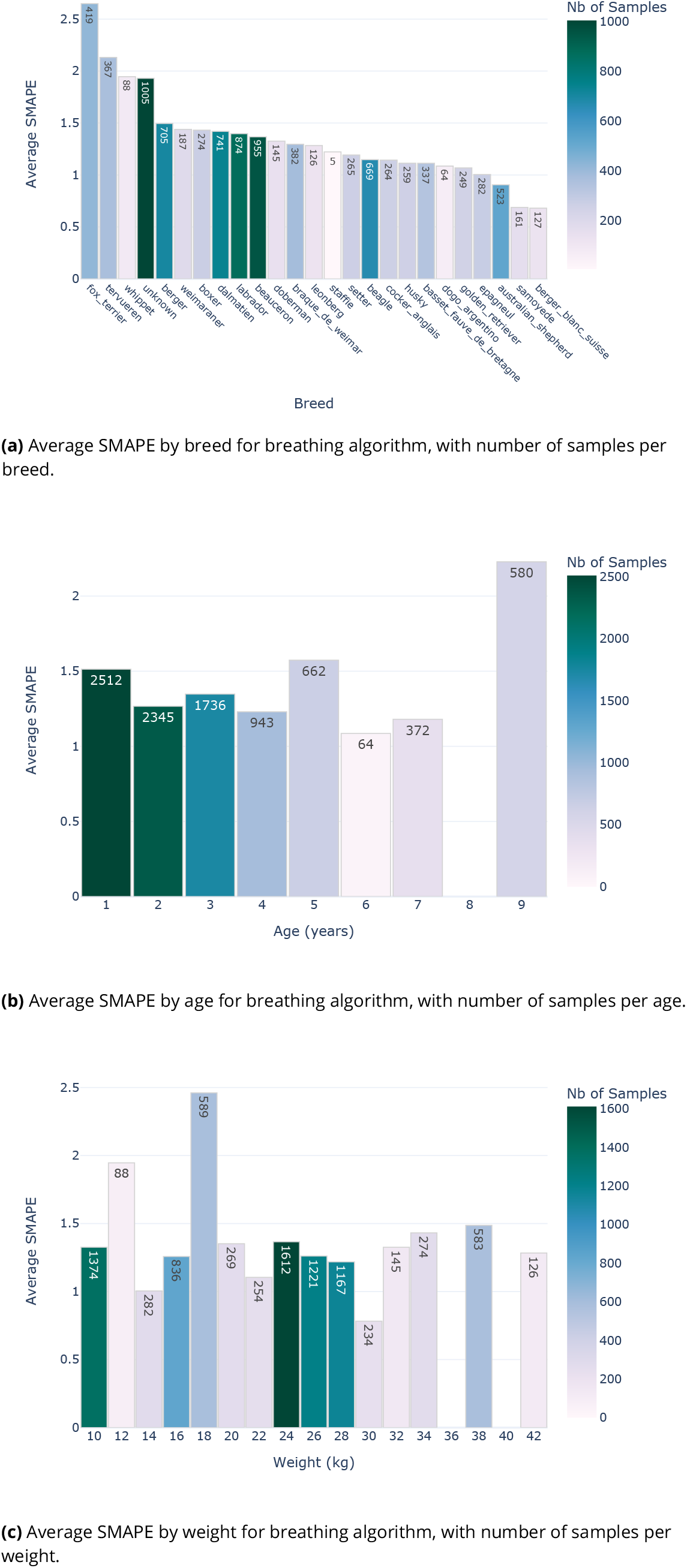
SMAPE distribution by breed, age and weight for breathing algorithm.

The average SMAPE is below 2.65% for all breeds in the test dataset, which show that the model is able to generalize well on all breeds. Similar results can be found for Age and Weight distribution, where the SMAPE distribution tend to be uniform across all ranges, with all bins having a SMAPE below 2.5%. These results show that our model is able to generalize well over every breed, age and size of dog, and could be used as a reliable tool to estimate the breathing rate of any dog on a daily basis.

We also give a more visual representation of the results by plotting the ground truth labels against the predicted labels, as shown on figure 14. As above, transparency is used to better show the density of points. We can see that the model is able to predict the ground truth labels with a high degree of accuracy, with a few outliers on some labels. These outliers are however not significant, as they represent a very small portion of the data. Indeed, when looking at the distributions of the total breathing rate for each dog, with the ground truth labels and the predicted labels, on figure 15, we can see that the distributions match closely for all dogs, with the exception of the dog 38, which is also the dog with the lowest number of samples. Therefore, its breathing rate distribution estimation may not be as accurate as the other dogs, due to a lack of validation data. We can see that all the dogs have a breathing rate within 5 to 35 MPM, wih a few outlying points.

**Figure 14.**
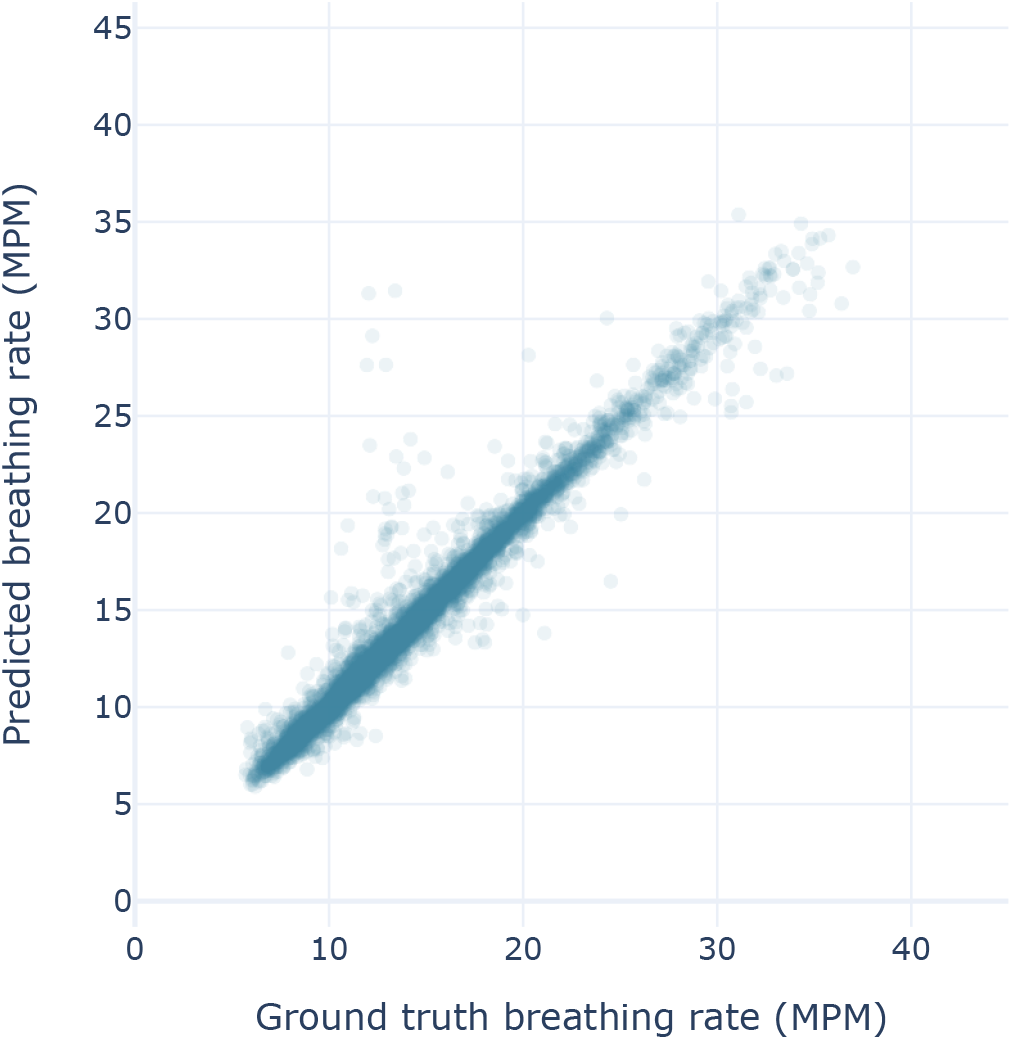
Ground truth labels against predicted labels for breathing algorithm. Transparency is used to better show the density of points. The diagonal represents perfect prediction.

**Figure 15.**
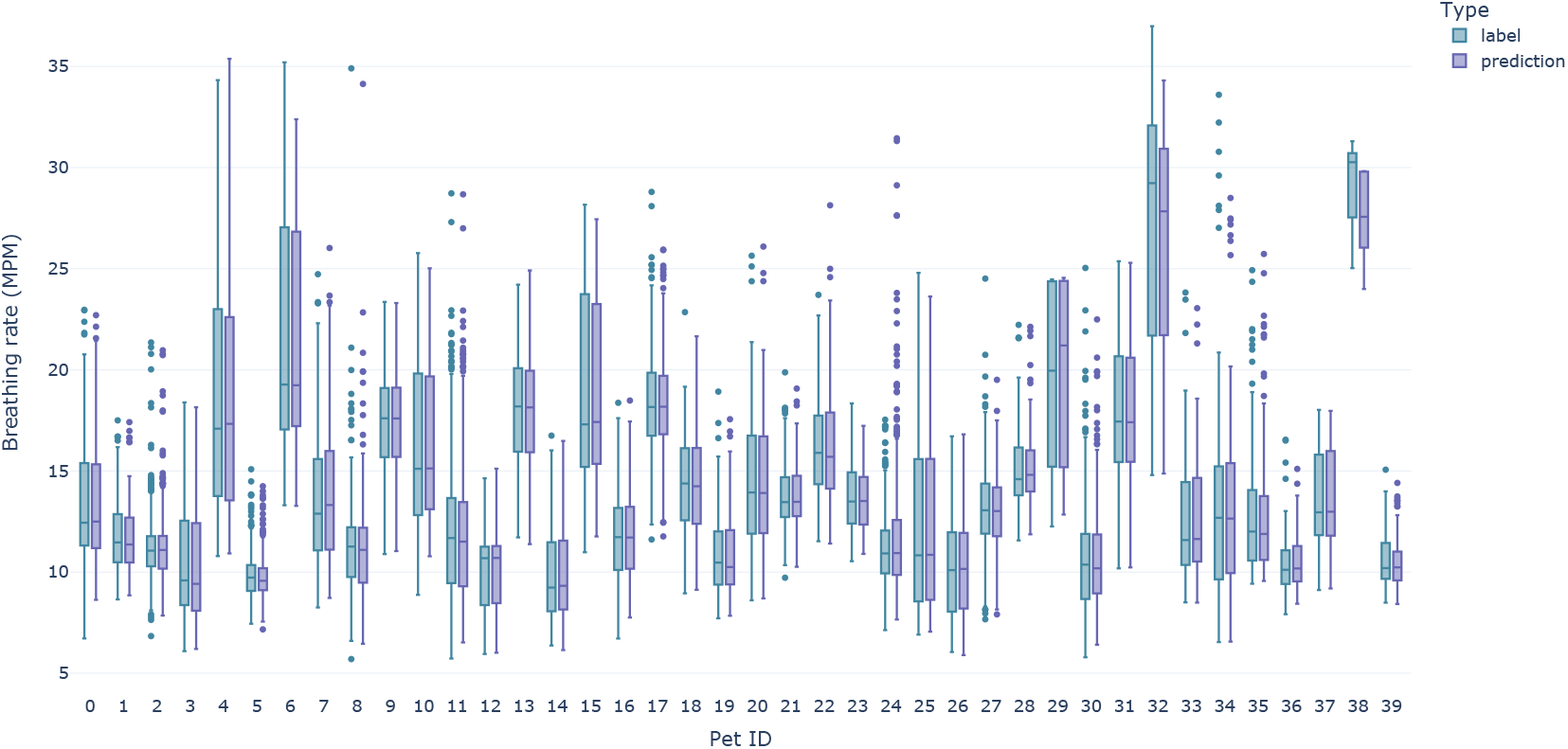
Breathing rate distribution (labels vs predicted) for each dog.

**Figure 16.**
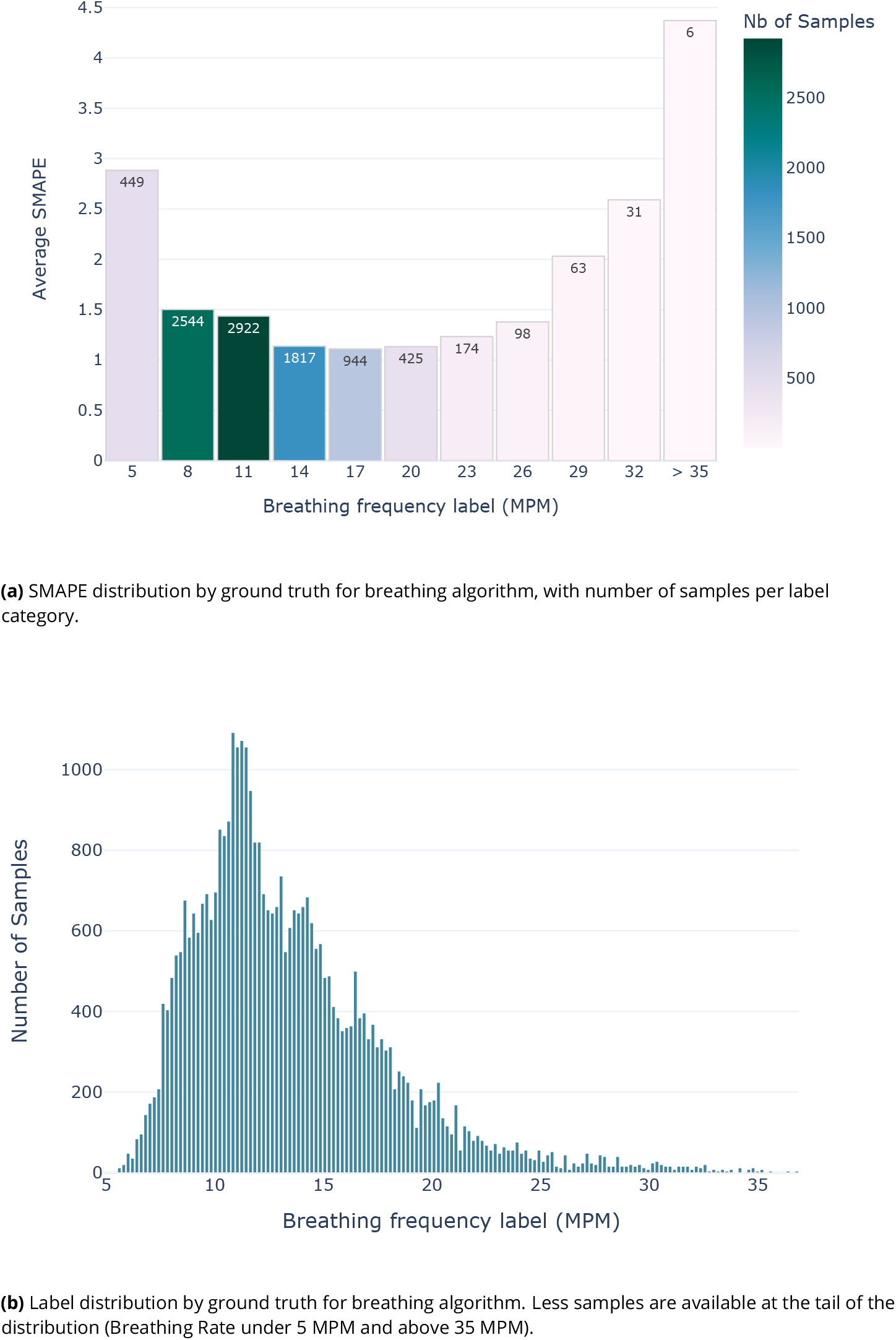
SMAPE and Label distribution by ground truth for breathing algorithm.

This distribution re?ects the current consensus on veterinarian medicine, which states that resting respiratory rates for dogs tend to fall on average between 5 MPM to 30 MPM (Porciello et al., 2016; Rishniw et al., 2012). This is clearly represented in our test labels distribution which approximates to a (skewed) normal distribution of mean 13.52 MPM with a standard deviation of 4.49 MPM. As more data will be collected with our collars, we hope to be able to gain more insight about the global distribution of breathing rates, especially on the lower and higher ends of the distribution.

### 5.3 Discussion

To the best of our knowledge, this is the first large-scale evaluation study to use a wearable device to measure the heart and breathing rate of dogs. With a representative dataset of 40 dogs, with a diverse range of breeds, ages and sizes, we show that our collar is able to accurately measure the heart and breathing rate of dogs, with a mean accuracy of 99.60% and 98.58% respectively, and is also able to detect heart beats with a mean F1 score of 98.04% at 50ms precision.

When compared to other studies, our collar provides a more accurate measurement of the dog’s heart and breathing rate, while being less invasive. Indeed, Brugarolas et al., 2015 and Foster et al., 2020a both use a harness to measure the breathing rate of the dog, which is more invasive than our collar. Compared to Foster et al., 2020b, our collar demonstrates a better breathing rate estimation accuracy (98.58% vs 94.3%) and a better heart rate estimation accuracy (99.60% vs 88.2%). Our f1score is also arguably better, although ours is computed at 50ms precision, versus 25ms for Foster et al., 2020b. Moreover, when compared to other human wearable devices (Jacobs et al., 2021, Li et al., 2019), our collar provides a more accurate measurement of the dog’s heart and breathing rate, with a mean absolute error of 0.44 BPM (±1.43 BPM) for heart rate and 0.38 MPM (±0.65 MPM) for breathing rate. For the breathing algorithm, 95% of our samples have an absolute error below 1.19 MPM, and 99% have an absolute error below 2.94 MPM, which is way below the acceptable limit of 5 MPM for a resting breathing rate measurement (Medical Instrumentation et al., 2002). Our breathing algorithm accuracy, at a 5 MPM precision, stands at 99.65%, which is also extremely good. For the heart algorithm, 95% of the samples have an absolute error below 3.82 BPM, and 99% have an absolute error below 6.92 BPM. The accuracy at 5 BPM is 98.8 %. Those results show that our collar has still good results even in the worst case scenario.

Overall, we argue that our algorithms perform very well on average, independently of the breed, weight, age and fur size of the dog. This is an interesting result, as it shows that the *Smart Dog Collar*’s algorithm is sufficiently general to work across all breeds and physical attributes, by only analyzing the small neck movements of the dog. These results support the hypothesis that our collar is precise enough to be used as a medical-grade tool to monitor the average resting heart and respiratory rates of the dog, on a daily basis, without the intervention of the dog owner.

## 6 Conclusion

Our results show that our collar is able to accurately monitor the resting heart and breathing rate of dogs, with an average SMAPE of 0.4% and 1.42% respectively (or 99.6% and 98.58% SMAPA) on the test data, regardless of the breed, age, weight or fur size of the dog. Our algorithms also outperform state-of-the-art devices, all the while being evaluated on a much larger dataset. Given the promising results obtained, the algorithm has been put in production and is currently being used to monitor the health of numerous dogs in the field. We are confident that it will be able to provide useful insights to dog owners and veterinarians alike by providing a continuous, non-invasive and objective measurement of the dog’s health. We also hope that the data collected can be used to further our understanding of the dog’s health such as the impact of stress or different cardiac pathologies on dog’s vital signs.

## Footnotes

1 This test dataset (containing ECG data and annotated labels) is made public on [Zenodo](https://zenodo.org/record/8020390).

2 This removed data is labelled as `bad_ecg` segments in our public dataset.

